# Cardiac myofibril networks induce shear stress

**DOI:** 10.1101/2025.11.10.687713

**Authors:** L. A. Murray, A. Quinn, C. Pinali., D.J. Collins, V. Rajagopal

## Abstract

Myofibril arrangement is critical to cardiac muscle function in health and disease. Historically, analysis of the impact of myofibril organisation on force and cell contraction has relied on the assumption of uniaxial arrays. However, improvements in imaging indicate that myofibrils form complex networks, though how these networks modulate force has yet to be explored. Here, morphological analysis of sheep left-ventricular cardiomyocytes is utilised to inform a non-linear finite element model of cell contraction. Analysis of deep learning segmentations of Z-Discs demonstrate that myofibrils are oriented about the major axis (mean = 0.03°) but deviate locally by up to 30° (standard deviation = 6.56°). Simulations produce unique deformations for geometries informed by myofibril orientations, displaying internal rotation and off-axis deformations. Moreover, anisotropy generates shear stresses distinct from the uniaxial case, demonstrating spatial relationships that balance shear across the cell. These findings highlight the impact of myofibril networks on forces during cell contraction.

## 1.0 Introduction

Cardiac muscle cells produce the forces underpinning each heartbeat. These forces are a product of synchronous cell-shortening, facilitated by organelles called myofibrils^1,2^. Myofibrils span the rod-like myocytes longitudinally in arrays, connecting 1.8-2 [μ*m*] contractile proteins, sarcomeres, which are attached in series and shorten up to 20% during activation^2,3^. Force is transmitted throughout the myofibril by Z-Discs which connect adjacent sarcomeric protein filaments, provide structural stability, and attach the myofibril to the cell membrane. Myofibrils force production, is facilitated by an influx of Ca^2+^ in a process called excitation-contraction coupling^2^. Ca^2+^ binds with proteins on the actin-filament within the sarcomere enabling interactions between actin and myosin filaments, producing shortening and decreasing the distance between adjacent Z-Discs^2,4^. Nevertheless, while the fundamental mechanisms of force generation has long been understood, there is increasing appreciation of the complexities and physiological implications resulting from the intracellular directions along which these forces are generated^5,6^.

Myofibril orientation has been historically presumed as uniaxial about the cell’s longitudinal axis^1,7,8^. These assumptions relied upon limited transmission electron microscopy (TEM) imaging, facilitating investigation in sarcomere structure and theories of contraction, but providing minimal 3D spatial context on global myofibril arrangement^7,9^. Moreover, early work utilised indirect-flight muscles which have been shown to be uniquely organised and uniaxial^7,10,11^. Therefore, intracellular stress has been historically decomposed into an *active* (axial) component produced by myofibrils, and *passive* (off-axial) components attributed to the cytoskeleton^12–14^. This assumption has been perpetuated in subsequent biophysical models^15–20^ and experimental studies^13,21–23^, though deeper examinations of myofibrils, particularly in mammalian samples, may provide a more detailed understanding of intracellular stresses and their relevance to cardiac physiology and disease conditions.

Importantly, volumetric EM imaging has demonstrated that myofibrils may display splits^11,24^, are asynchronously aligned^25,26^, and form interconnected branches^6,10^. Goldspink originally highlighted myofibril structure in mice bicep brachii, observing that distal sarcomeres ‘split’ into smaller ‘daughter’ sarcomere along the myofibril length^8^, later suggesting splitting acts as a modulator of tension development or proliferation in fast- and slow-twitch muscles^24,27,27^. These splitting events were also observed by Tomanek in kitten skeletal muscle^28^. More recently, Jayasinghe et al. applied laser scanning confocal miscopy to study Z-Discs as an indicator of myofibril structure, observing that myofibrils are helictical over myocyte length^25^. The images of rat, rabbit, and human cardiomyocytes similarly display an asynchronous alignment of Z-Discs, purporting that these structures may positively influence contraction synchrony. Further, with focused ion beam-scanning electron microscopy, Willingham et al. highlighted that myofibril networks form in fast-, slow-twitch, and cardiac myocytes through sarcomere branching events^6^. The result was recreated by Ajayi et al.^10^ through selective gene-knockout of *Drosophila* myocytes, both groups hypothesising an impact on myocyte force production. Nevertheless, how such multiaxially aligned myofibrils impact intracellular stress has not been investigated.

An understanding of stress dynamics within the cell is critical for appreciating function in health, disease, and exercise. Cardiac myocytes respond to their mechanical environment, modulating membrane channels, calcium signalling, and force transmission^29–31^. These regulatory behaviours may help myocytes respond to altered loading conditions in disease states^32^, as well as facilitate development^33,34^, and accommodate demand^15^. Understanding intracellular stresses is key to understanding the mechanosensitive behaviours of key organelles such as mitochondria, which have been shown to deform during contraction^35^ and modulate calcium levels^36^, and the nucleus, whose migration is impacted via myofibril force^34^. Nevertheless, while experimental designs incorporate off-axis stress behaviour^37,38^, virtually all biophysical models assume myofibrils produce zero shear stress.

In this work we examine the intracellular stresses in a striated myocyte resulting from off-axis Z-Disk orientations. This process is highlighted visually in Fig. 1. Here, serial-block face scanning electron microscopy (SBF-SEM) images of sheep left ventricular myocytes were segmented to create a physiologically informed biophysical model. A U-Net++^39^ was trained on manual segmentations to inform morphological analysis of Z-Discs as a proxy for measuring myofibril orientations. This analysis indicates that myofibrils permute at a distribution of angles within the network about the contraction axis (Fig. 1, red arrow; *μ* = 0.03°, *σ* = 6.56°). Angles were then interpolated into a hyper-elastic Finite Element (FE) model using the FEniCSx^40–43^ open-source package^40^. The FE model is parameterised to fit existing experimental trends for myocyte tension^12,17,44,45^. Integration of myofibril multiaxial alignment results in shear behaviours that are not reproducible with a uniaxial assumption. Taken together, this study indicates that incorporating multiaxially alignment in biophysical models is vital for describing intracellular stresses in cardiac myocytes.

**Fig 1.**
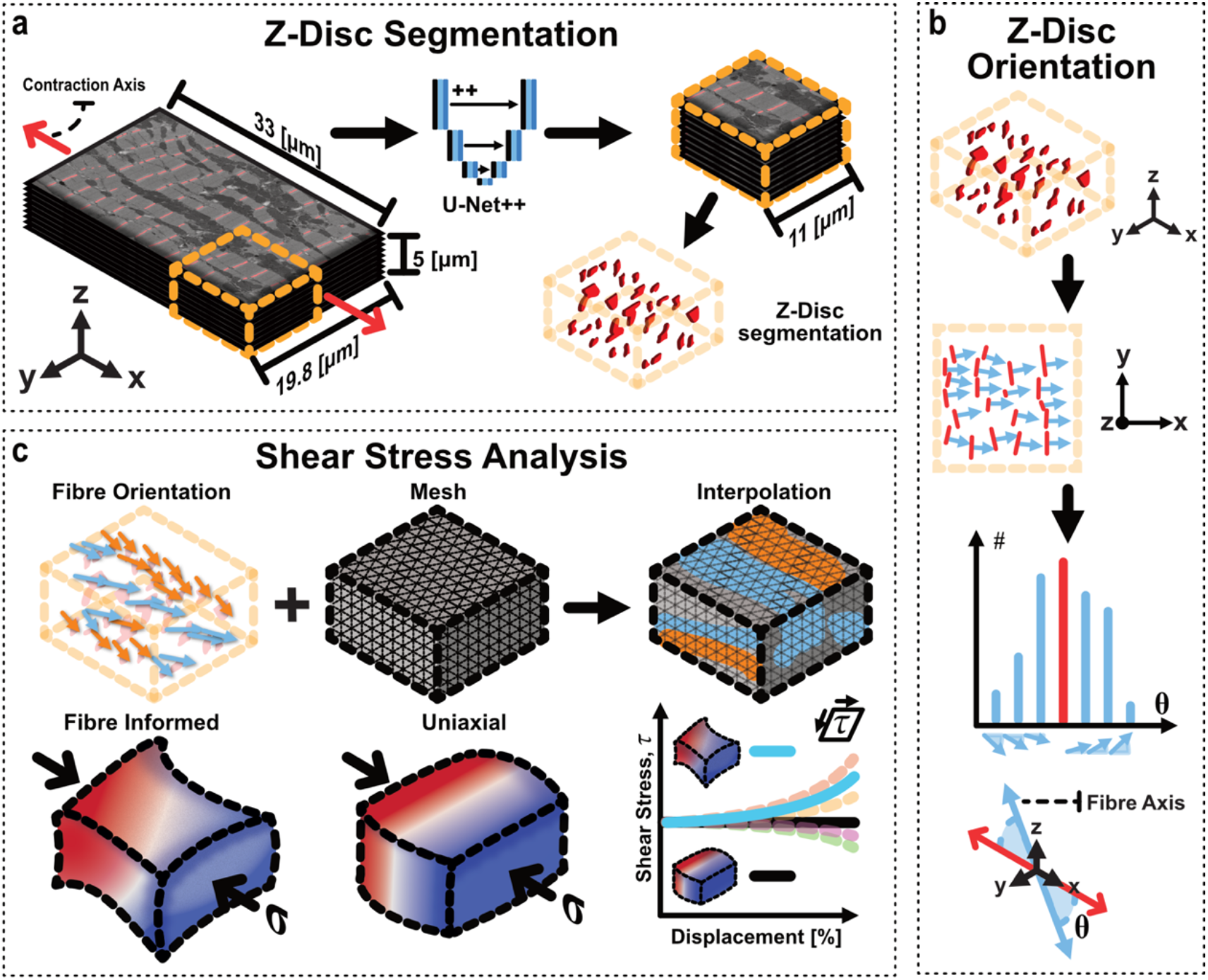
Visual abstract demonstrating the workflow from cardiomyocyte segmentation through to morphological analysis and mechanical simulation. (a) Z-Discs are isolated from portion of sheep left-ventricular cardiomyocyte (**33** × **19. 8** × **15 μm**; one **5 *μm*** volume indicated here) through U-Net++ Machine Learning model instance segmentation. (b) Isolated segmentations of Z-Disc are analysed with principal component analysis to quantify orientation of fibre in reference to contraction axis of cell. (c) Fibre orientation interpolated into non-linear mechanical model for contraction simulation. Fibre-informed simulations and uniaxial tests are contracted and shear stress quantified. Quantification indicates that shear stress in fibre-informed cases produce a distribution not reproduced by uniaxial model.

## 2.0 Results

### 2.1 Z-Disc segmentation reveals myofibrils are not uniaxially aligned

Isolated Z-Discs create varied patterns that do not predominantly align with the contraction axis. The larger 3D cell volume is partitioned into smaller volumes and regions (Fig. 2a, green, red and blue). The bottom layer (Fig. 2b) is characterised by a large longitudinal grouping of mitochondria that span between edges, interrupted only by a nucleus on the proximal end. The boundaries of this mitochondria region and the nucleus contort the neighbouring sarcomeres’ orientation and shape, similar to results shown elsewhere^46^. This structure is more pronounced in the top-down 2D render shown at the bottom of Fig. 2b, where the Z-Disc axial alignment is clearly impacted by the mitochondria and the nucleus.

**Fig 2.**
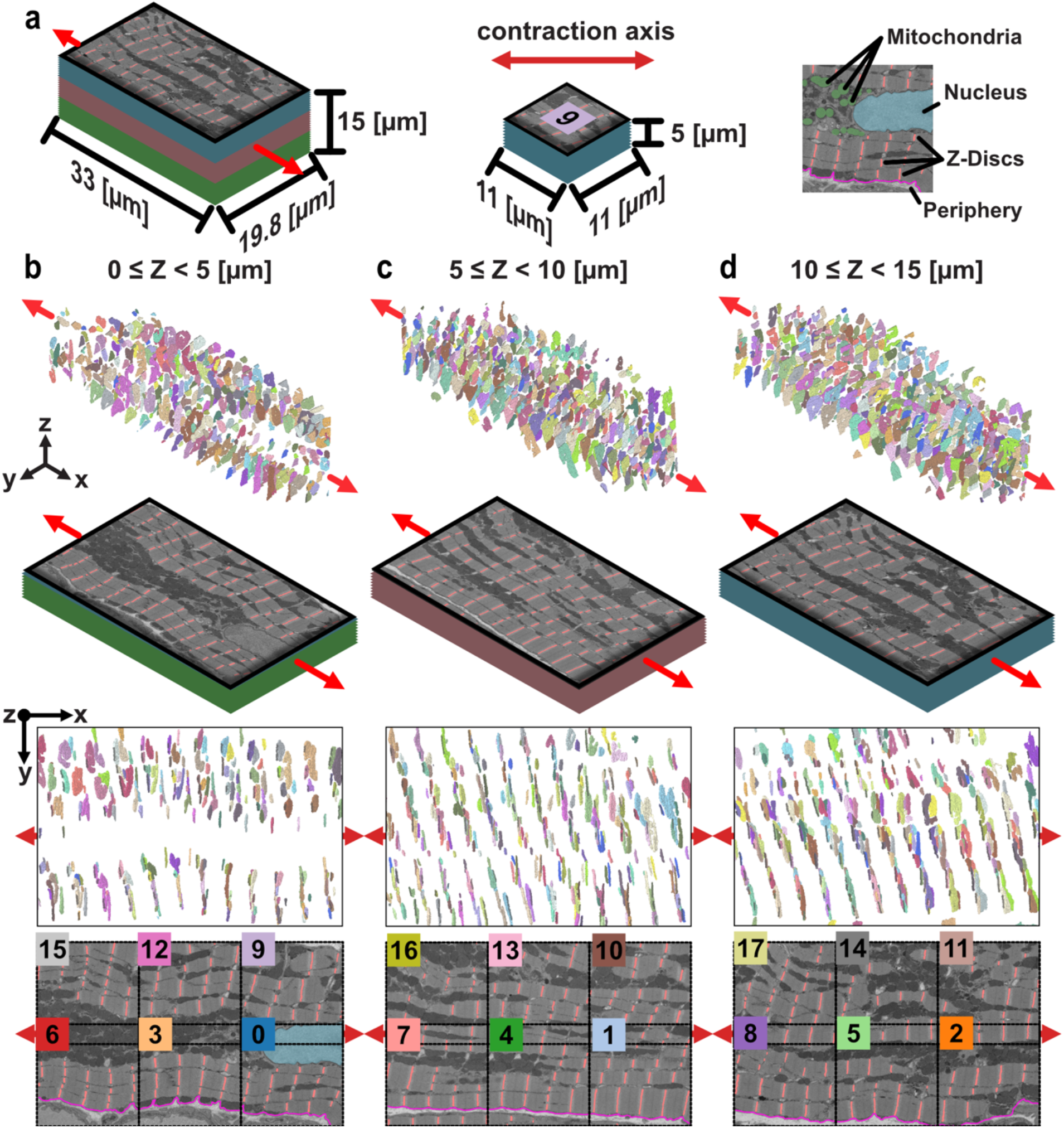
Z-Disc segmentation from cardiomyocyte volume and split into test regions. (a) Top left, key indicating dimensions of full cell segment (**33** × **19. 8** × **15** [***μm***]), broken into three stacks. Middle, dimensions of test regions (**11** × **11** × **5** [***μm***]) displayed. Right, extraction from Region 0 with key cell landmarks indicated; mitochondria in green, nucleus in light blue, Z-Discs in red, and periphery in pink. (b-d) Visualisation of Z-Disc instance segmentations, electron microscopy images, and test regions for (b) **0** ≤ ***z*** < **5** [***μm***], (c) **5** ≤ ***z*** < **10** [***μm***], and (d) 1**0** ≤ ***z*** < **15** [***μm***]. (b-d) Row 1 (top), Orthogonal render of Z-Disc segmentations with contraction axis indicated. Row 2, Orthogonal view of EM slices. Row 3, 2D top-down view of Z-Disc segmentations aligned left-to-right with contraction axis. Row 4 (bottom), 2D top-down view of EM slices with overlay of numbers of regions for identification; dimensions as indicated in (a).

In each volume the regular ∼2 *μm* spaced Z-Discs along myofibrils is apparent (see Fig 2b-d bottom row), although Z-Discs are not vertically aligned over width (see Supplementary Material 1). Asymmetry in alignment between myofibrils has recently been explored^25^, however the Z-Discs in this sample only exhibit alignment across width within myofibrils bundles, with a loss of this coherence between myofibrils that are spatially separated. Z-Discs do not appear aligned over the full cell width.

Differences in myofibrillar arrangement patterns are apparent near the cell’s lateral periphery and nuclei. 2D visualisation of the slices (Fig. 2b-d bottom) highlight the Z-Discs at the cell periphery (all slices) and about the nuclei (Fig. 2b bottom). The cell periphery aligns myofibrils with the cell contraction axis, with a reduction of orientation changes particularly apparent in the second volume (Fig. 2c). Investigation into Z-Discs surrounding the nuclei, however, show a contouring of bodies about its border (Fig. 2b bottom). In both cases statistical testing did not indicate any significant differences in orientations about these cell landmarks (see Supplementary Material 2).

### 2.2 Myofibrils vary as much as 30° from longitudinal axis

Principal component analysis (PCA) of segmented Z-Discs indicates that myofibrils throughout the cell are oriented between −30° and 30° of the contraction axis (Fig. 3; standard deviation *σ* = 6.56°; mean *μ* = 0.03°), which was calculated from the cell’s long axis (see Methods). To compute this angle the isolated Z-Discs were processed to determine the *normal* vector (Fig. 3a) through the third principal component. The Z-Disc is directly connected to the *actin-* and *titin-* filaments which run perpendicular to the Z-Disc body and parallel with contraction^1^. The normal vector is then compared to the contractile axis of the cell to determine spherical deviation (see Fig. 3a middle illustration).

**Fig 3.**
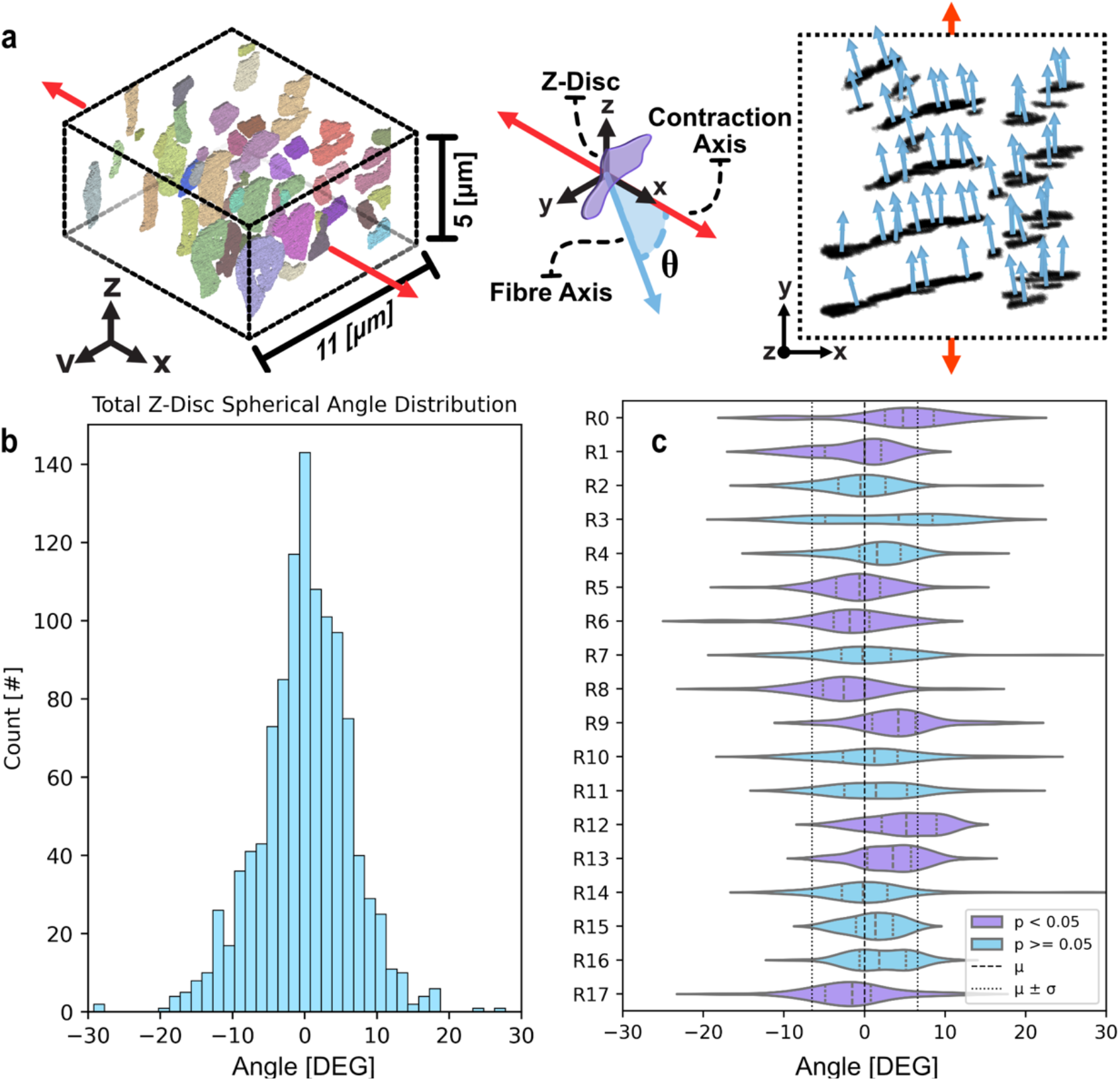
Isolation of Spherical Angle distributions from Principal Components Analysis of segmentation data. (a) Top left, Region 7 rendered in orthogonal view. Top middle, diagrammatic indication of relationship between Z-Disc, Fibre-Axis, and Contraction Axis; the spherical angle is computed through the vector angle between the Z-Disc normal vector and contraction axis. Top right, 2D top-down view of PCA computed Normal vectors to segmentations from Region 7 shown top left. (b) Histogram of global orientation values for all segmented Z-Discs, bin size: **1. 5**°. (c) Violin plots with quartiles shown for orientation data per test region. Purple regions indicate statistical significance between regional orientation values and global set. Significance was determined with Man Witney U-test and Welch’s t-test ***α*** = **0. 05**. Vertical dash (--) and dotted (:) lines indicating global quartiles.

Myofibrils are oriented symmetrically about the major axis (*n* = 1138, *skew* = −0.16) with a standard deviation (*σ*) of 6.56° and mean (*μ*) of 0.03° (see Fig. 2b). The large kurtosis value (*κ* = 3.49) of this distribution indicates a concentration of values within the first standard deviation. Indeed, whilst a uniaxial model of myofibrils presumes complete alignment, only ∼12% of instances had an orientation within 1° of the major longitudinal axis.

Regional orientations exhibit significant deviations in distribution and mean with respect to the main axis. Compared to the global set with Mann Whitney U-test and Welch’s t-test, 9 of the 18 test sets demonstrated significance (see Fig. 3c; see Supplementary Material 2). Whilst region size impacts statistical relevance, these distributions nevertheless highlight the regional variability in myofibril arrangement. Further, of regions exhibiting a statistically significant orientation distribution, all but one (R17) were immediately adjacent to another significant region. Of the 10 significant regions, 5 included the cell periphery and 2 included the nucelus, though these landmarks did not significantly impact adjacent orientations when tested independently with the same Mann Whitney U-test and Welch’s t-tests (see Supplementary Material 2).

The median of myofibril orientation also shifts across the cell volume, where Fig. 4a and 4c indicate that fibre orientation is heterogenous. Interestingly, slicing through the *x*-axis (Fig. 4c, left) depicts a transition from a positive deflection to a negative deflection. This observation mimics Fig. 3c, where Regions 0-8 and Regions 9-17 exhibit a decreasing trend in median orientation, the former being a larger gradient. The largest median positive orientations (R0, R9, R12 – see Fig. 3c) all appear closer to the distal end (0 ≤ *X* < 11 [*μm*]) of the cell, whereas the largest negative median orientations all occur on the proximal end (22 ≤ *X* ≤ 33 [*μm*]; R6, R8, R17). Observing slices through the *y*-axis (Fig. 4c, middle) confirms this behaviour. Therefore, the cell proceeds from positive- to negative- median orientation shift over its length.

**Fig 4.**
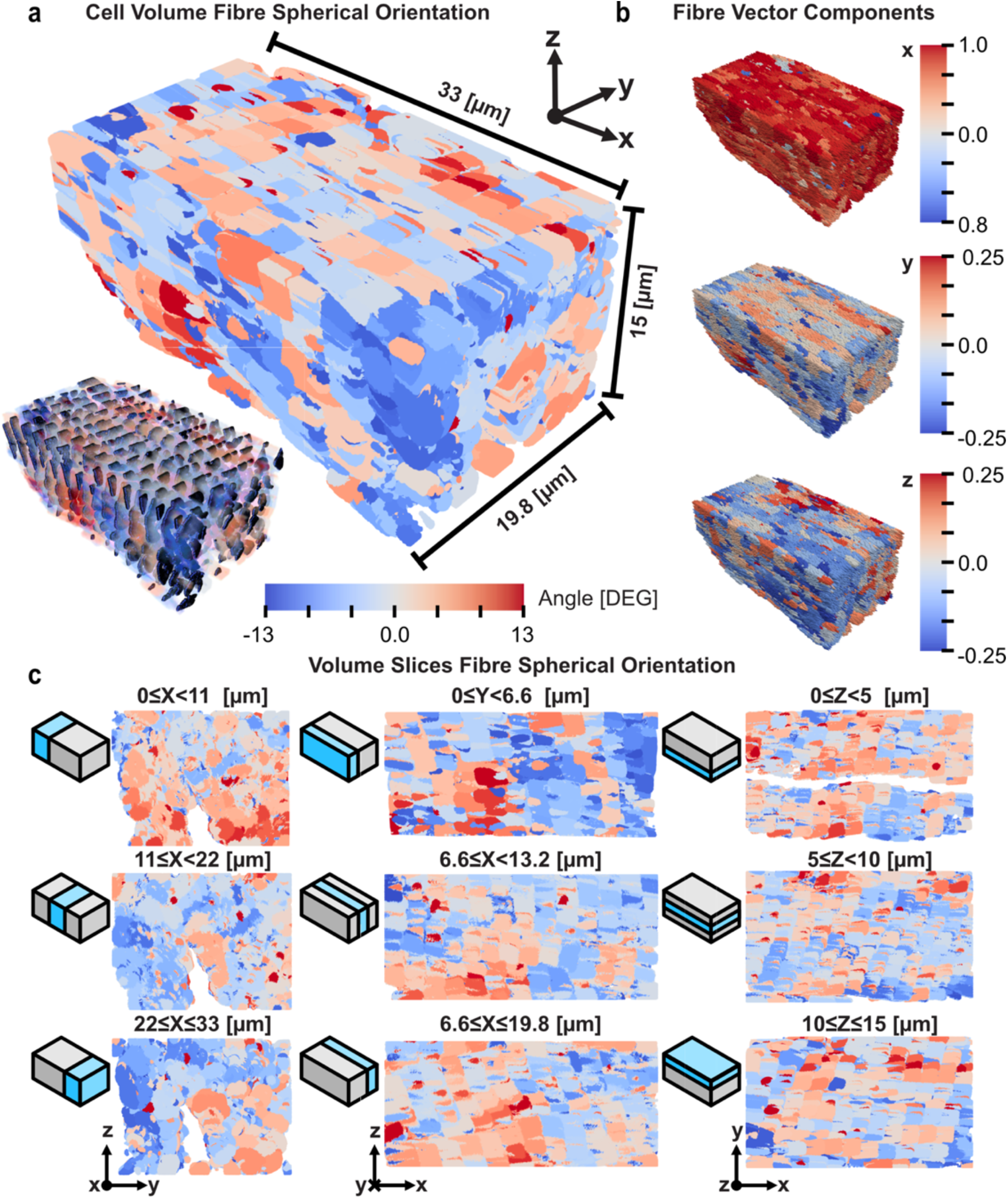
Interpolation of fibre orientation into cell volume. (a) Whole cell volume render with interpolation of Z-Disc orientations throughout lengths of sarcomere sections. Colour spectrum is limited to two standard deviations of global orientation values. Overlay of Z-Disc segmentations onto orientation render displayed in lower left. (b) Glyph render of cartesian vector components for each Z-Disc from top-to-bottom: ***x***-component, ***y***-component, ***z*** -component. ***x***-component demonstrates homogeneity in principal direction as anticipated, ***y*** and ***z*** show larger variability. (c) Axial slices through the rendered volume. Each slice is a third of the dimension’s length. Note void in ***x***-slices (left) which correspond with the large mitochondria group and nucleus observable in top-right ***z***-slice. Slicing passes through Z-Discs and sarcomere unevenly, therefore Z-Disc patterning is less obvious.

### 2.3 Myofibril anisotropy introduces uneven deformation and rotation

A computational model of myofibril organisation and large nonlinear deformation mechanics was implemented using FEniCSx^40–43^ to investigate the influence of myofibril organisations on muscle contraction. Myofibril orientation was introduced into simulation meshes with an interpolation of orientation into mesh nodes. A tensor push-forward transformation then allowed for development of metric tensors and the Christoffel symbol of the second kind, with

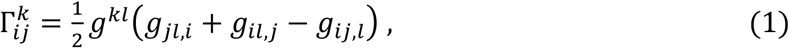

where *g*^*ij*^ and *g*_*ij*_ are deformed contravariant and covariant metric tensors, and the Christoffel symbol is necessary for adjusting derivatives over the deformed volume with the covariant derivative,

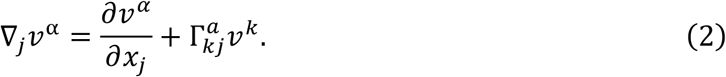

The covariant derivative forms a necessary part of variational form of the biophysical model with incompressibility restraint

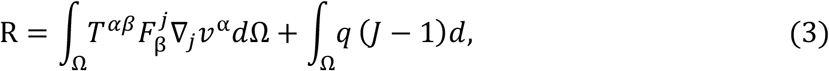

where *T* is the second Piola-Kirchhoff tensor, *F* is the deformation gradient tensor and *J* is the third invariant. Contraction was simulated with stepwise displacement boundary conditions on the maximal and minimal faces of the mesh.

Despite rigid displacement boundary conditions and incompressibility, myofibril anisotropy introduced asymmetrical and region-specific deformation. Comparison of the boundary *y* - displacement across mesh segments with the uniaxial deformation indicates that all test regions increase variance in deformation alongside a median shift (see Fig. 5a-b).

**Fig 5.**
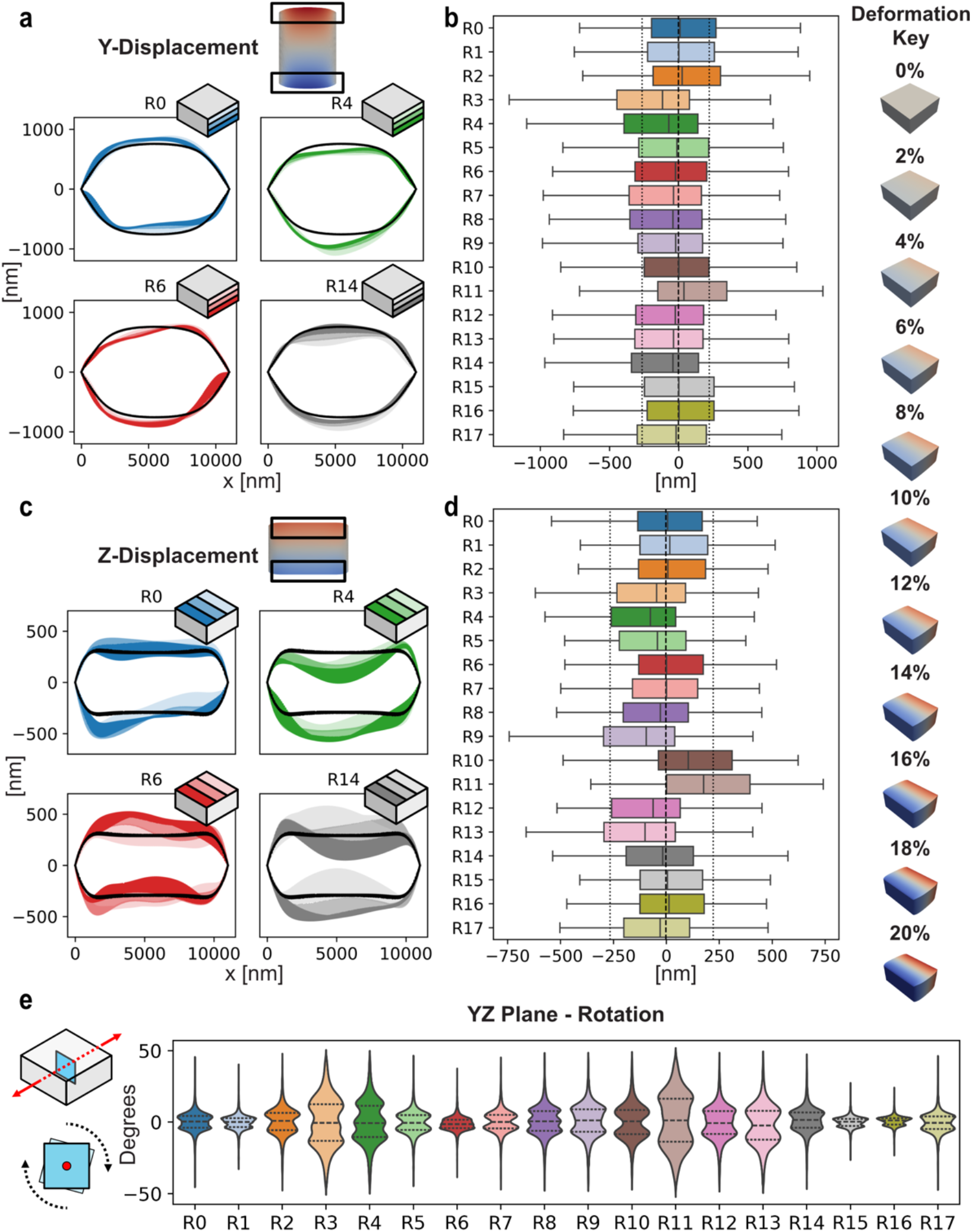
Region displacements and in-plane rotation at 20% displacement. (a-b) ***y***-displacement visualized as select region shaded boundaries (a) and all regions boxplot distributions (b). (a) Boundary displacements for Region’s **0**-**4** are overlayed onto displacements of uniaxial test. Displacements are plotted for thirds of the dimension to illustrate transition of deformation over length. Darkest shade indicates first slice, closest to origin (***y*** = **0**). (b) Boxplots of each Regions displacement values throughout volume. Dashed (--) and dotted (:) lines indicate quartiles of uniaxial test. (c-d) ***z***-displacements visualized to similar capacity as (a-b). (e) ***yz***-plane rotation produced by displacement displayed as violinplots with quartiles. Plane rotation determined at inner plane to reduce impacts of rigid boundaries. Violin plots show the distribution of common rotation values for each region, see Region 4 indicating frequent rotation values at −**25**°, **0, 25**°.

Four regions were randomly selected to more closely investigate how boundaries deform with anisotropy. Fig. 5a and 5c plot the simulation mesh boundaries on *y*- and *z*-faces overlayed onto the boundaries of the uniaxial case (Fig. 5a, 5c; black solid lines). The simulation boundaries are displayed with three shaded sections depicting the nodal displacement in thirds of the volume due to their rapid changes over the surface. These regions depict how anisotropy deflects the volume’s maximum and minimum boundary irregularly in both the *y*- and *z*-axis. z-displacement in Regions 6 and 14 further indicate that despite a minimal median shift from the uniaxial case (Fig. 5d) there is notable deviation across the region’s profile; R14 begins with a predominantly negative deflection before progressing positively (see shading in Fig. 5c bottom right). Therefore, despite volume constraints, introducing anisotropy creates boundary displacement deflections.

Myofibril anisotropy further introduces internal rotation. Measurement of rotation about the screw axis in the *YZ*-plane in our physiological sample exhibits unique profiles for all test cases (Fig. 5e). In contrast, the uniaxial case produces 0° of rotation in the YZ-plane or across streamlines in the displacement field (see Supplementary Material 3). Internal rotation varies predominantly between −25° and +25°, increasing with a larger median shift in displacement; direct comparison of scale of violin distribution (Fig. 5e) matches median shifting in displacement boxplots (Fig. 5d; for example, Regions 3,4,9,10, and 11). Whereas more uniformly displaced tests that align medially with the uniaxial example have demonstrate less rotation (i.e. Regions 6 and 16). Larger changes in deformation profiles due to anisotropy result in increased rotation od nodes within the mesh (see Supplementary Material 3).

### 2.4 Anisotropy creates shear stress and unique spatial profiles

Anisotropic myofibrils produce unique shear stresses not reproducible by a uniaxial model. The Cauchy stress tensor (Fig. 6a; diagrammatic), calculated through conversion from the first Piola Kirchoff tensor (see Eq. 5), is decomposed and plotted for each displacement component in Fig. 6b.

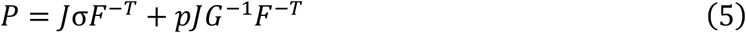

**Fig 6.**
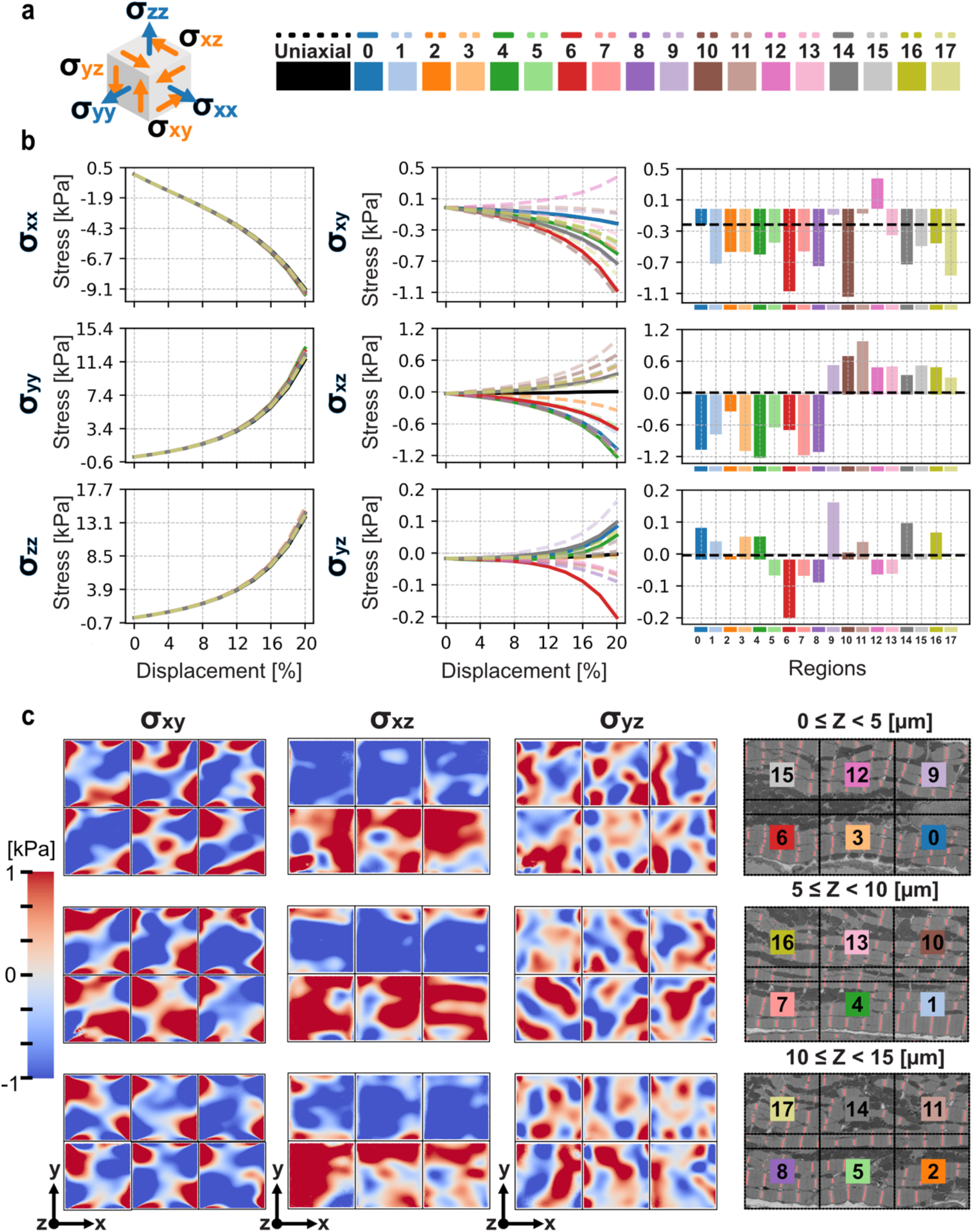
Stress trends and spatial shear for simulations at 20%. (a) Left, diagrammatic visualization of stress tensor components. Top-right, render of uniaxial deformation for each simulation step change, bottom-right, test label key for stress trends and bar charts. (b) Left, normal stress components plotted over displacement; nearly all test cases show minimal deflection from the uniaxial test, this is likely an artifact of rigid boundary displacement. Shear stress trend (middle) and barcharts (right) for each test condition. Shear trends demonstrate variance in shear profile as deformation reaches 20%. Barcharts are ordered numerically with region numbers, dashed (--) line indicates uniaxial test value. Values shown are final deformation stresses at 20%. (c) Spatial shear profiles for each region test case aligned with EM images of slices. All distributions are displayed with the same colour bar range. Some pattern is evident in ***xz***-shear which corresponds with barchart (b, middle).

The average normal stress components in each case are similar in response to the identical displacement boundary conditions for each simulation (Fig. 6b, left). However, all anisotropic simulations produce unique shear responses (Fig. 6b, middle and right). These profiles are most pronounced after 12% displacement, suggesting that anisotropy may be less relevant for early stages of contraction. Full contraction^3^, 20% strain (Fig. 6b right), demonstrates variability between all cases. Clearly, anisotropic regions, even with minimal orientation mean-shift, produce shear stress.

Shear stress in the *xz*- and *yz*-components is distributed between positive and negative inflection for simulations (Fig. 6b, right). In contrast, *xy*-shear was decidedly negative for all but one test, R12, which had a large positive skew in orientation distribution (see Fig 3c); *xy*-shear was also the only non-zero shear for the uniaxial case. This non-zero shear for uniaxial simulation result is likely due to the artificial degree of freedom constraint on the *x*-axis faces. Normalising for the uniaxial case results in four simulations with positive stress, though this may still be artificially constrained. *xz*- and *yz*-shear, however, produce null shear in the uniaxial case with notable distribution ratios of positive and negative shear stress for all simulations (*xz*: 1: 1, *yz*: 8:6, *positive*: *negative* for tests with noticeable shear).

Shear stress exhibits a spatial relationship over the cell volume like that observed in the orientation behaviour. The balance of shear stress displayed by the *xz*-component, visualized by the bar charts in Fig. 6b, corresponds with the spatial shear distributions (Fig. 6c). *σ*_01_ is noticeably negative for all upper regions of the cell slices (Fig. 6c left), with a positive value for all lower slices. This effect is most apparent for Regions 0-2 and 9-11 which oppose sides of a nucleus.

## 3.0 Discussion

This study reveals that cardiac muscle cell myofibrils create a distribution of orientations and produce unique intracellular shear stress distributions. This finding challenges long-standing assumptions based on a uniaxial model, which has treated muscle contraction as occurring purely under axial loading since Huxley’s foundational work in the early 1900s^7^, as well as more contemporary ultrastructural imaging work^1^, biophysical simulations of fibre force^16,20^, and experimental measurements^12,47^.

Understanding the production of shear stresses has the potential to elucidate the impacts of multi-axial force trajectories, where existing biophysical models have so far limited active stress dynamics to uniaxial forces. However, lateral force transmission is an important feature of musculoskeletal physics^18,48^ and has burgeoning appreciation in cardiac constitutive models^49,50^. Recent work has demonstrated that lateral forces change in disease states^51^ and theorise its role in inter-cellular transmission^48,52^. While historical models of myofibril arrangement do not characterise these dynamics, our physiologically informed biophysical model demonstrates the presence and magnitude of shear stresses that would contribute to multi-axial force.

Visualisation of the spatial shear stress relationship, and observations of internal rotation, indicates that the cell experiences internal torsion during contraction. As displayed in Fig. 5e, all simulations experienced internal rotation due to anisotropy. Similarly, in Fig. 6c shear in the *xz*-component indicates that lateral sides of the cell experience opposing shear. This finding indicates that anisotropy produces internal rotations during contraction, and more broadly that shear stress may produce torsion about the cell’s major axis. Moreover, coupling of adjacent regions would further quantify the impact of shearing across the volume.

Reports have demonstrated that muscle cell architecture is impacted by fibre type^6,26^. Cardiomyocytes require a constant provision of energy dissimilar from skeletal muscle, impacting the size, orientation, and connectivity of mitochondria and sarcomeres^26^. Similarly, Willingham et al.^6^ displayed that branching events in myofibrils are increased in slow-twitch muscles compared to cardiac and fast-twitch variants, whereas Ajayi et al.^10^ indicated that myofibril branching is impacted by gene knock-out in *Drosophila*. This work demonstrates that non-axial orientation produces shear stresses significantly different from a uniaxial model in a physiological model of a cardiomyocyte, future work, therefore, may focus on how other muscle cell architectures impact shear stress.

The hyperplastic anisotropic biophysical model presented here is a foundational model which has the further potential to incorporate other cell physiology. Active contraction and calcium dynamics, for instance, are important contributions that have been incorporated into other multi-dimensional cell models^19,20,53^; incorporation of these data could be used to produce a more detailed physiological model. Further, changes to the rigid boundary conditions considered here may impact how extreme anisotropy diverts from the uniaxial model.

## 4.0 Conclusion

In this study we investigated, to our knowledge, the largest and most detailed Z-Disc segmentation of a cardiomyocyte and produced the first fibre-orientation informed non-linear myocyte deformation model. Deep learning segmentation of the cardiac cell confirmed that Z-Discs produce a distribution of orientations values concentrated about the major axis. These values varied up to 30° and were not significantly impacted by organelles. However, fibre orientation did display characteristic differences across the cell geometry with a gradient of positive to negative mean-shift over the cell length.

Myofibril anisotropy produces shear stresses that are not reproducible with the uniaxial model. Whilst normal stresses were maintained, physiologically informed simulations created arrays of shear values. All simulations produced shear components of greater magnitude than the uniaxial case, the latter resulting in no shear in the *xz*- and *yz*-components. This study indicates that incorporating off-axis myofibril orientation is necessary to account for shear stresses produced during cardiac muscle cell contraction.

## Methods

### Z-Disc Segmentation

The ultrastructure of sheep left ventricular cardiomyocytes were captured with scanning-block-face electron microscopy (SBF-EM) as previously described^54^. Within this volume (6000 × 6000 × 359 [*pixels*]; *X and* 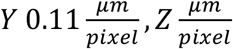), three cardiomyocytes were present and 19 random regions (1024 × 1024 × 100 [*pixels*]) were isolated for manual annotation of Z-Discs in Napari^55^. These annotations were then used to automatically segment the largest present cardiomyocyte. The chosen cardiomyocyte was a full cell depth and half-length of similar cardiomyocytes (∼94 [*μm*])^56^.

A smaller initial subset of these patches were annotated and used to train a 2D U-Net++^39^ with a ResNet-50^57^ weights using unified focal loss^58^. The output segmentations from this original model were used to create a segmentation on the remaining patches which were manually corrected before creating the full set to train the final model. During each epoch smaller patches from the 19 annotated 3D blocks were extracted to introduce training set variability. Sampled 2D patches were 384 × 384 pixel covering 70% area, resulting in 1,648 patches per epoch.

Output segmentation voxels were generated through 2D smoothing tiling of overlapping predictions. Manual corrections were made to erroneous segmentations after inspection and before utilization in further analysis. In total 1138 Z-Discs were segmented and included in this study, with manual annotation regions comprising ∼ 14.9% of the whole volume and automatic segmentation comprising ∼59.9%.

### Morphological Analysis

For morphological analysis orientations were normalised to the cell’s contraction axis. This was defined as the long axis of the cell and validated later by the average orientation angle of Z-Discs (see Fig. 3). From here, instances of Z-Discs were analysed to inform on centroid position, pixel-size, and orientation. Centroid data, pixel quantities, size, and elliptical axis were calculated with the scikit-image^59^ *measure* module.

Fibre orientation was defined through the Z-Discs’ normal vector in direction of cell’s major axis. To determine the normal vector *Principal Components Analysis* (PCA)^60^ was utilised; the first two components characterizing the structure of the segmentation, as the major and minor axis of length and width. The third component is, by definition, perpendicular to the first two, and normal to the face. PCA was implemented with scikit-learn^61^.

The spherical orientation was calculated as the angle between the vectors of the third principal component and contraction axis (see Fig. 3a). To achieve this all third components were required to be pointing in the positive x-axis. The resulting distribution of values is displayed in Fig. 3b with a bin size of 1.5°. Bin sizes larger than this are insufficient for appreciating variance, smaller sizes increase noise.

### Mesh

A prism mesh defined on the dimensions of cell regions was created in Gmsh^62^. The geometry (11000 × 11000 × 5000 [*nm*]) was defined with second-order tetrahedral elements. Mesh refinement was tested to ensure convergence of solution and minimization of adjacent node difference. The resulting test mesh had 358,656 elements and 463,120 nodes.

### Constitutive Equation

The Guccione type constitutive equation (see Eq. 1) for orthotropic behaviour was implemented^50^.

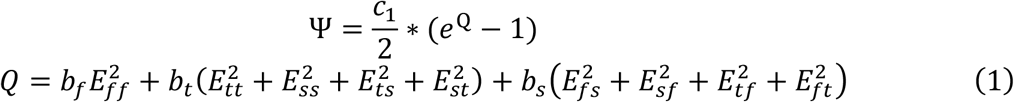

Here, *b*_*t*_ and *b*_*s*_ were set as equal reducing the equation to transverse isotropic as previously explored^14,63^. The remaining constants (*b*_*f*_, *b*_*t*_, *c*_1_) were optimised to approach a maximum stress value of 9 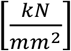 (or [*kPa*]); balancing the range of experimental measurements of cardiac tension^12,44,53^. Solutions converged to *b*_*f*_ ≈ 12.68 [*kPa*], *b*_*t*_ ≈ 11.04 [*kPa*], *c*_1_ ≈ 2.81.

### Regions Orientation Interpolation

Myofibril orientation was interpolated into the mesh by first mapping the nodal positions of the mesh to the Z-disc pixel data from segmentation regions. This was supported with SciPy’s ^64^ KD-tree. Once mapped, angle data was interpolated into second-order (10-node) tetrahedral Lagrange functions with the in-built FEniCSx^40–43^ architecture. Nodes within the Z-Disc boundaries were assigned to the same angles as those which were directly mapped to the nearest-neighbour tree.

To ensure continuity between Z-Discs, Gaussian smoothing was applied strictly over Z-Disc nodes and the sarcomere regions between. Smoothing boundaries were based on the diameters of Z-Discs (disc-axis) and length of sarcomeres (fibre-axis). Reports suggest cardiac Z-Discs range from 100 [*nm*] to 140 [*nm*] ^65,66^, and sarcomere slack length of 1.8 − 2 [*μm*] ^1,3,67^; these values corresponded to the standard deviations provided to the smoothing function.

### Fibre Field

Anisotropy was incorporated into the variational calculus similarly to Nash and Hunter^68^ and Guccione et al.^50^. First, undeformed basis vectors (*A*_*i*_) and metric tensors (*G*_*ij*_) are calculated via rotation of cartesian coordinates to orient with the myofibril data in each region. Here, *G*_*ij*_ is the inner product of the basis vectors,

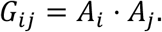

This is interpolated into Quadrature points within native FEniCSx^40–43^. FEniCSx utilizes the Lagrangian form of variational calculus, therefore the Euclidean metric covariant and contravariant tensors are also determined. Lower case *g*_*ij*_ and *a*_*i*_ are used as convention to indicate deformed tensors.

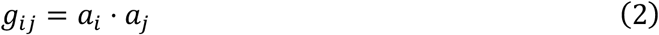

Subsequentially, the Christoffel Symbols of the second kind,

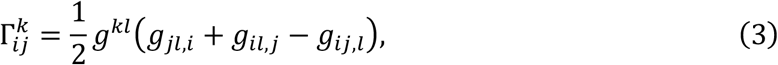

and covariant derivatives,

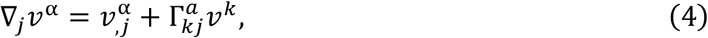

are calculated to update the variational form

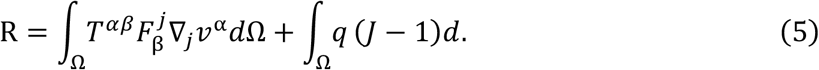

The First-Piola Kirchoff is calculated per a push-forward transform on the Cauchy stress tensor.

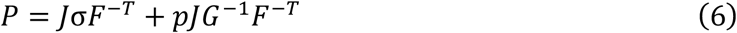

### Boundary Conditions and Simulation

Contraction was simulated with discrete displacement of the regions at maximum and minimum *x*-position. The planes were fixed in *y*- and *z*-axes with the *x*-axis displaced 20% (2.2 *μm*) to achieve a percentage displacement. All other degrees of freedom move freely.

An iterative non-linear solver native to FEniCSx^40–43^ was employed. The solution was incremented at 2% intervals. Convergence was determined with an incremental tolerance calculation, requiring a tolerance of 10^.;^ and a maximum of 50 iterations per Newton Solver. Simulations were run on the University’s High Performance Computer and were allocated 512GB per node and run serially to avoid node-mismatch in parallel.

### Displacement and Rotation

Boundary displacements for each region were compared by taking the edges of the deformed shape. To compare *z*-Displacement, values along *x*-axis were plotted for three regions through the *y*-axis. Maximum and minimum displaced values on the upper and lower edge of these regions were overlayed (see Fig. 5). Similarly, *Y*-Displacement was displayed across the *x*-axis for three regions across the *z*-axis.

Rotation data were extracted from the centre region and calculated in the *Y*-*Z* plane about *x*-axis. Two vectors were produced once each datapoint was centralized to the origin: the first being a vector to the undeformed point, the second being to the deformed point. The angle difference between the two with the *y*-axis was provided as the angle of rotation.

### Stress and Strain Calculation

Stress values were extracted from an internal 90% volume. This bounding allowed for the reduction of boundary artefact in the solutions and overestimations of stress. Cauchy stress was calculated with the inversion of Eq. 6 yet omitting hydrostatic pressure. Hydrostatic pressure was interpolated with first-order (4-node) Lagrange tetrahedra.

Each tensor component was then calculated by taking the mean stress over the internal volume in each direction. The stresses displayed in Fig. 6 correspond with the principal directions and relevant shear terms. All spatial visualisations were achieved in Paraview^69^.

## Supplementary Material 1

**Fig S1.**
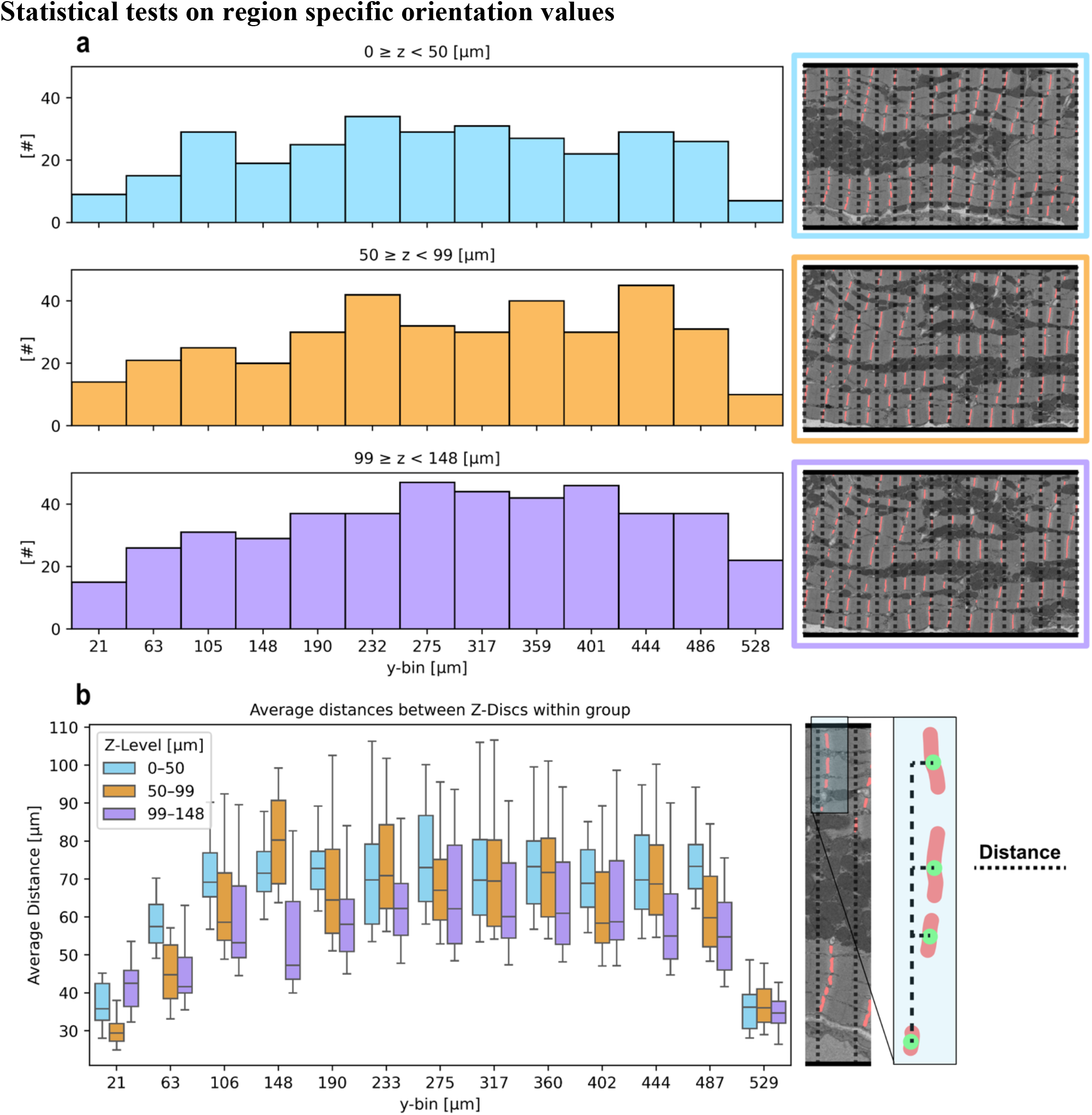
Distances between grouped Z-Discs across cell width. (a, left) Histograms of Z-Discs centroids across three difference z-axis slices. (right) Images of Z-Disc regions indicative of corresponding histogram; overlay of dashed (--) lines to represent boarder of bins in histograms. Distributions show similarly of shape as Z-Discs across cell fall within bins. Each slice demonstrates a similar quantity of Z-Discs. (b, left) Boxplots of inter-bin distances between each centroid (see right for indication) and those within the same group. Smaller groups have tighter distribution of distances due to decreased sample sizes. Boxplot spread indicates Z-Discs centroids are not aligned over cell width.

## Supplementary Material 2

**Fig S2.**
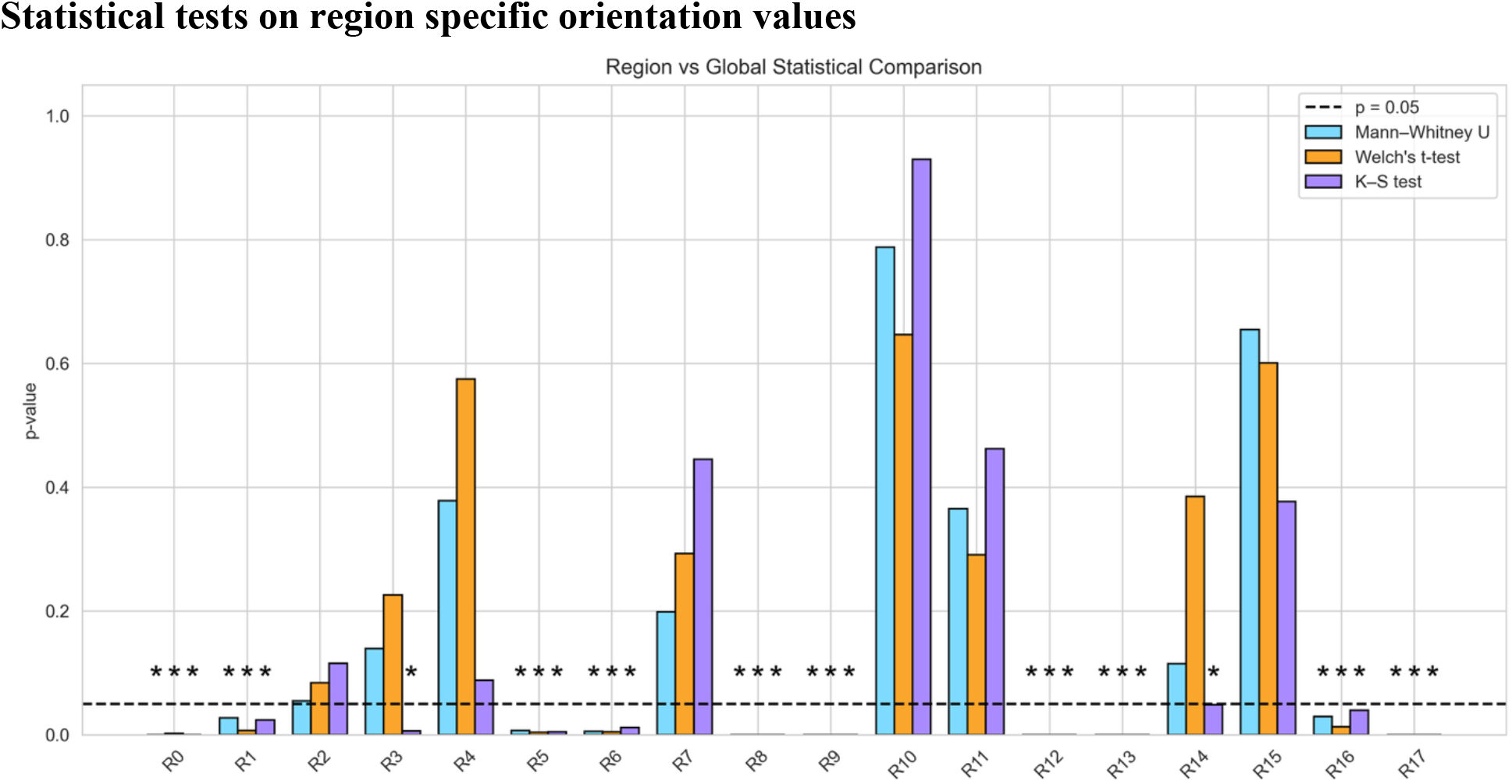
Barplot of p-values from statistical tests on region orientation data. p-values displayed for Mann-Whitney U-test (light blue), Welch’s t-test (orange), K-S test (purple), per test regions. Horizontal dashed (--) line indicating an ***α*** of **5**% (***p*** − ***value*** = **0. 05**). Stars (*) used to indicate p-values below the threshold *p<0.05, **p<0.01, ***p<0.001.

**Table S1.**
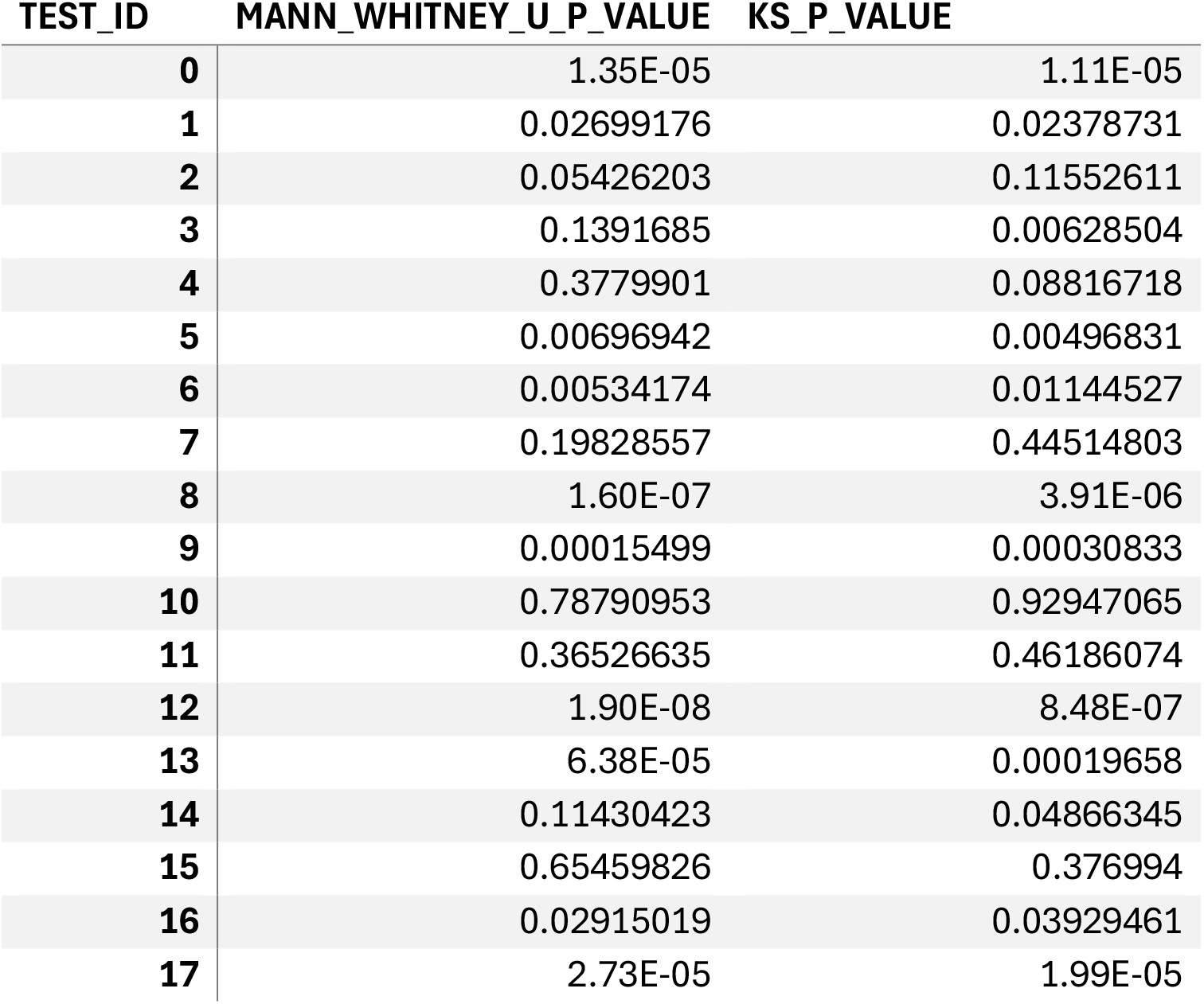
P-values for statistical tests.

| TEST_ID | MANN_WHITNEY_U_P_VALUE | KS_P_VALUE |
| --- | --- | --- |
| 0 | 1.35E-05 | 1.11E-05 |
| 1 | 0.02699176 | 0.02378731 |
| 2 | 0.05426203 | 0.11552611 |
| 3 | 0.1391685 | 0.00628504 |
| 4 | 0.3779901 | 0.08816718 |
| 5 | 0.00696942 | 0.00496831 |
| 6 | 0.00534174 | 0.01144527 |
| 7 | 0.19828557 | 0.44514803 |
| 8 | 1.60E-07 | 3.91E-06 |
| 9 | 0.00015499 | 0.00030833 |
| 10 | 0.78790953 | 0.92947065 |
| 11 | 0.36526635 | 0.46186074 |
| 12 | 1.90E-08 | 8.48E-07 |
| 13 | 6.38E-05 | 0.00019658 |
| 14 | 0.11430423 | 0.04866345 |
| 15 | 0.65459826 | 0.376994 |
| 16 | 0.02915019 | 0.03929461 |
| 17 | 2.73E-05 | 1.99E-05 |

**Fig S3.**
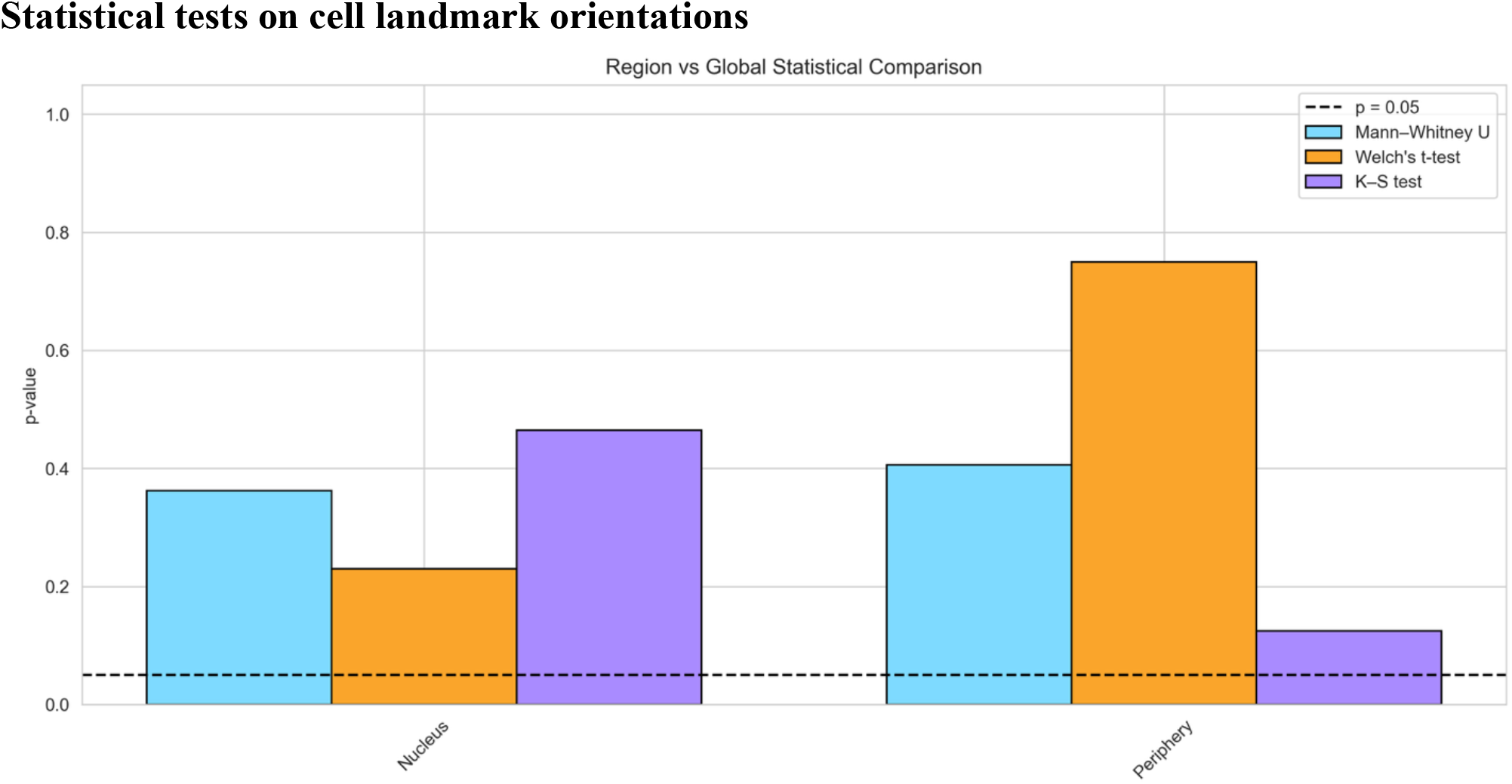
Barplot of p-values from statistical tests on cell landmark orientations. p-values displayed for Mann-Whitney U-test (light blue), Welch’s t-test (orange), K-S test (purple), per test regions. Horizontal dashed (--) line indicating an ***α*** of **5**% (***p*** − ***value*** = **0. 05**). Stars (*) used to indicate p-values below the threshold.

**Fig S4.**
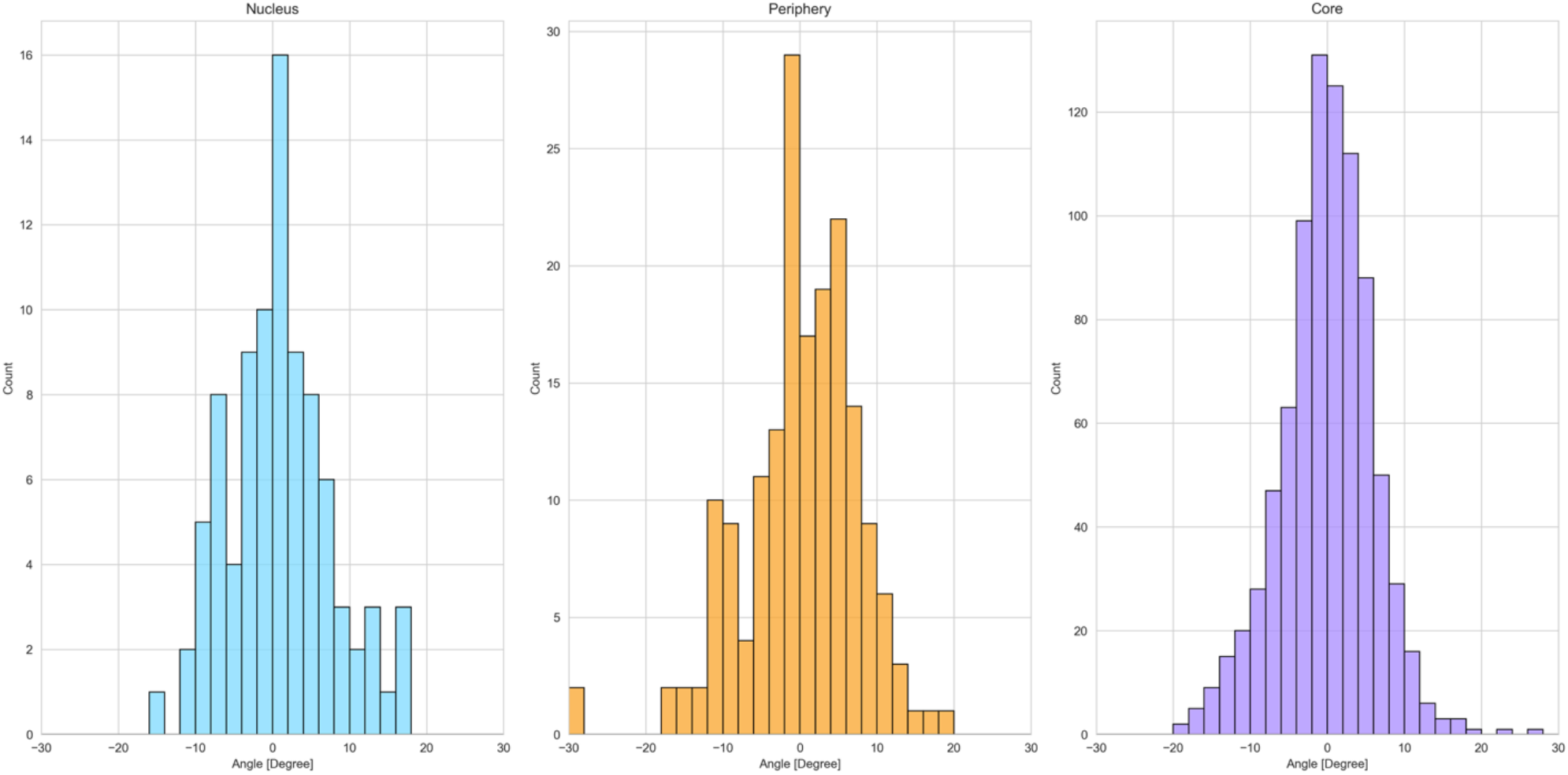
Histograms of landmarks orientation Z-Disc values. Distributions for nucleus (left; blue), periphery (middle, orange), and core (right, purple) regions. Domain is between −**30**° and **30**°. Z-Discs directly adjacent to landmarks are included.

## Supplementary Material 3

**Fig S5.**
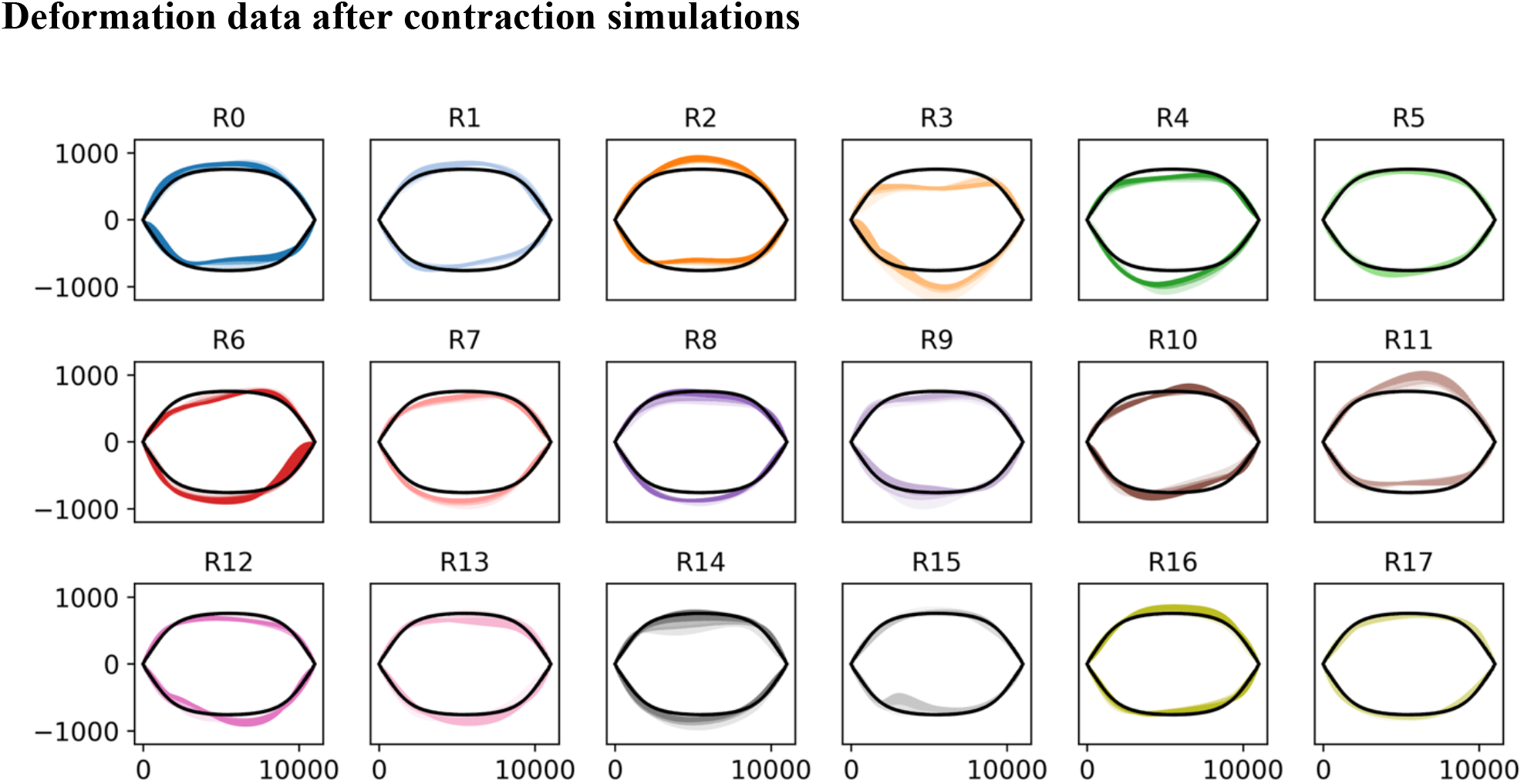
Y-Displacement of boundaries for each simulation overlayed on uniaxial case. as in Fig. 5] Black solid lines are the boundary surface of the uniaxial simulation. Each coloured simulation is broken into three shaded components indicating thirds of the boundary depth.

**Fig S6.**
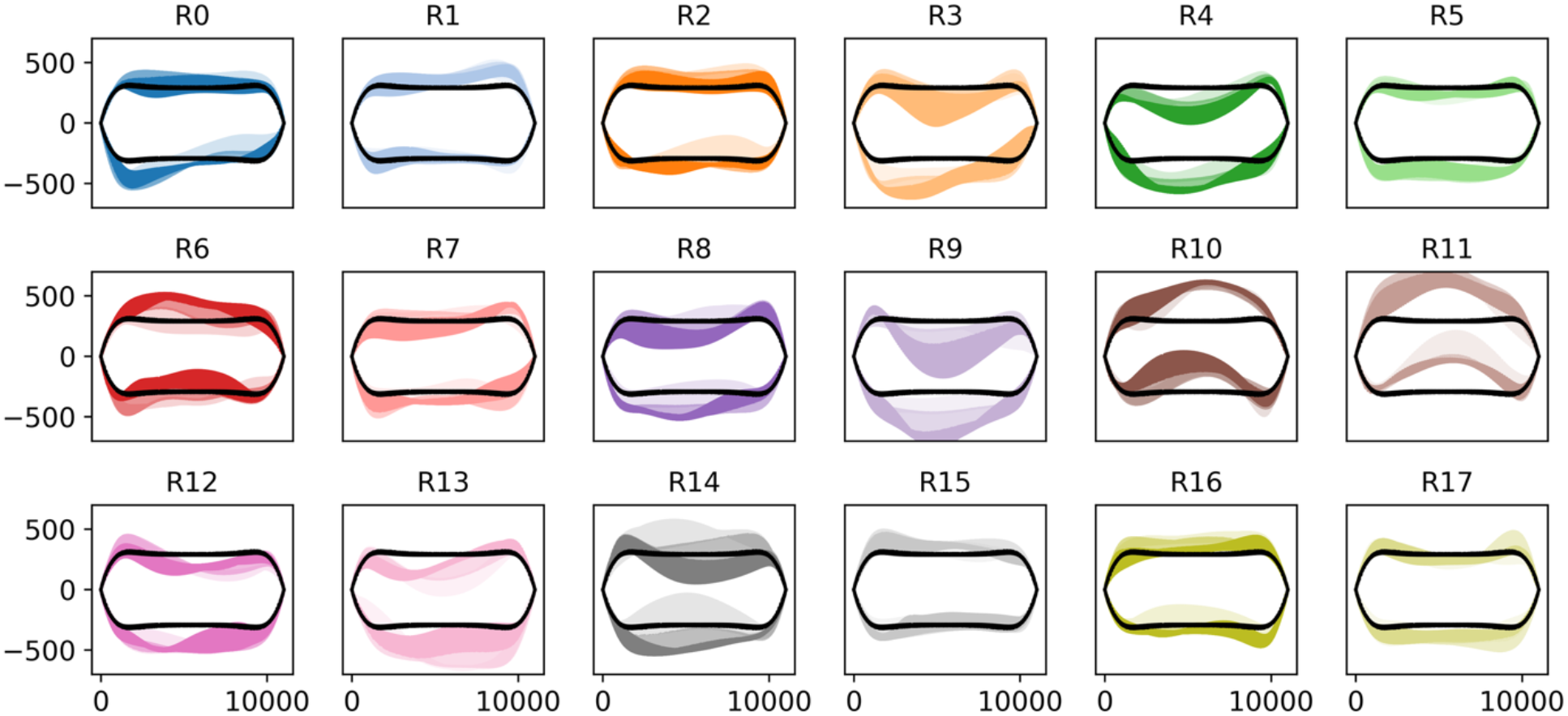
Z-Displacement of boundaries for each simulation overlayed on uniaxial case. [as in Fig. 5] Black solid lines are the boundary surface of the uniaxial simulation. Each coloured simulation is broken into three shaded components indicating thirds of the boundary depth.

**Fig S7.**
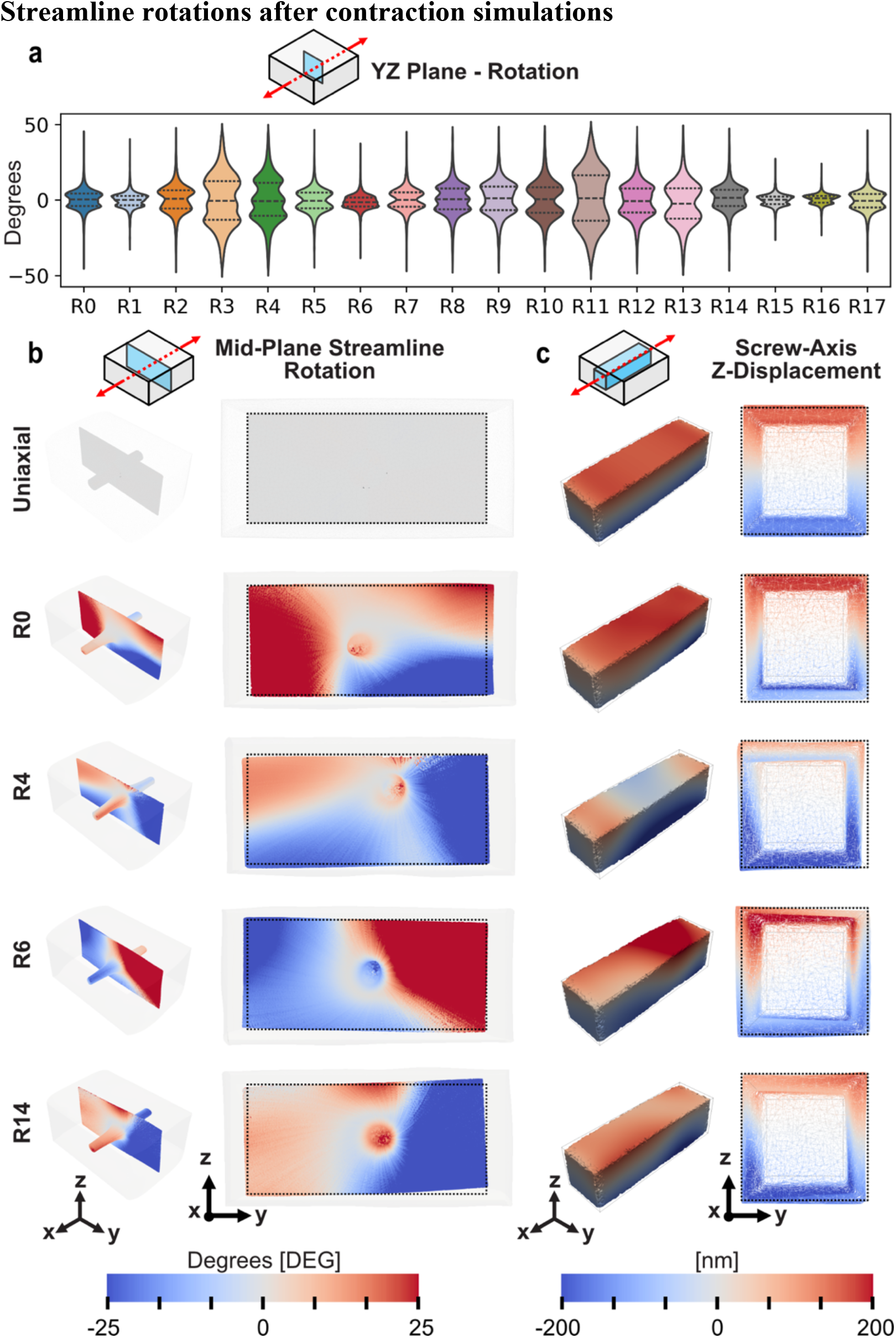
Streamline rotations in select simulations with internal displacement. (a) [as in Fig. 5] ***yz***-plane rotation caused by displacement displayed as violinplots with quartiles. Plane rotation determined at inner plane to reduce impacts of rigid boundaries. Violin plots show the distribution of common rotation values for each region, see Region 4 indicating frequent rotation values at ≈ −**25**°, **0, 25**°. (b) Streamline plots of mid-plane rotation for select simulations and uniaxial case. Streamlines move from the centre of each test and merge in mid-plane. Streamline follows displacement gradient and is inevitably uniform for uniaxial contraction. (c) Displacements of internal beam about screw axis displaying unique behaviour in test cases due to internal rotation.

